# Mapping 3D Heterogeneity of Thyroid Tumors Using Micro-CT based Radiomics

**DOI:** 10.1101/2025.06.13.658854

**Authors:** Kiarash Tajbakhsh, Olga Stanowska, Marija Buljan, Antonia Neels, Aurel Perren, Robert Zboray

## Abstract

Tumor heterogeneity plays a central role in treatment resistance, disease progression, and diagnostic uncertainty. However, it may be overlooked by traditional 2D histology. Accurate 3D assessment of tumor microarchitecture is therefore essential for capturing its spatial complexity. Micro-CT, an emerging imaging modality that offers high-resolution 3D virtual histology of soft tissues, provides a promising alternative. Combined with radiomics—a computational approach for interpretable quantification of tissue phenotypes—this technique enables a deeper understanding of tumor biology beyond visual inspection. In this study, we analyzed a large cohort of thyroid tumors (418 patients) using micro-CT imaging of next-generation tissue microarrays, from which we extracted radiomics features. We achieved robust classification of (i) neoplastic versus non-neoplastic thyroid tissues, (ii) papillary versus follicular thyroid carcinoma, and (iii) BRAF V600E mutation status. Feature interpretation using Shapley additive explanations revealed key visual traits driving these classification decisions. Preliminary results also indicated radiomic patterns associated with TERT promoter mutations, suggesting the existence of potential surrogate imaging biomarkers. Overall, micro-CT radiomics shows strong potential as a complementary tool for improving diagnostic and prognostic accuracy in thyroid cancer and offers a novel platform for quantitative pathology into the 3D spatial complexity of neoplastic tissues.

## 1 Introduction

Tumors are dynamic entities that evolve following a Darwinian model of clonal selection and adaptation. Environmental and therapeutic pressures can convert an initially monoclonal lesion into an ecosystem whose subclones differ in molecular profile, morphology, and stromal context. Some of these subclonies can develop drug resistance, replicative immortality, and cooperative metastasis [1, 2]. Despite this complexity, thyroid malignancies are still classified primarily by conventional histology, which may overlook critical spatial heterogeneity and thus limit current diagnostic frameworks.

Five major histotypes of thyroid cancer are recognised—papillary (PTC), follicular (FTC), poorly differentiated (PDTC), medullary and anaplastic carcinoma—with PTC and FTC accounting for 80% and 15% of cases, respectively [3, 4]. Prognostic assessment in PTC primarily relies on architectural and cytological features, whereas the diagnosis and risk stratification of FTC depend on the presence and extent of capsular and vascular invasion. While mitotic activity and necrosis are important prognostic indicators in both subtypes.

Borderline tumors blur these criteria: the follicular variant of PTC (FVPTC) shows both PTC- and FTC-like characteristics and suffers from particularly low inter-observer agreement [5, 6]. Furthermore, the focal nature of capsular invasion in FTC renders conventional histology vulnerable to sampling bias, increasing the risk of both over- and under-diagnosis [7]. This challenge is further compounded by interobserver discordance regarding what constitutes significant vascular invasion.

Molecular profiling is becoming increasingly important in the diagnosis and prognostics of thyroid carcinomas [8]. In PTC, co-occurrence of BRAF V600E and TERT-promoter mutations identifies tumors with aggressive trajectories; in FTC, TERT-promoter mutations and RAS alterations likewise portend poorer outcomes [9, 10, 11]. Crucially, these driver events are themselves heterogeneous: TERT mutations can be patchy within FTC [12], while multifocal or polyclonal PTCs may harbor discordant BRAF and RAS genotypes [13, 14, 15]. Such evolution-driven intratumoral heterogeneity complicates diagnosis, risk stratification, and ultimately precision therapy.

Current diagnostic methods fall short in capturing the three-dimensional architecture and spatial distribution of tumor heterogeneity. PCR-based TERT assays, for example, remain time-intensive and costly, limiting their clinical utility. Immunohistochemical tests for RAS only recognise a subset of point mutations. The histomorphological and molecular variability guides treatment decisions, and are all based on 2D histology and tumour-wide molecular testing. The distribution of molecular changes in 3D setting is unknown. Consequently, an innovative diagnostic tool that integrates both 3D structural information and molecular data would aid in evaluation and improve the management of thyroid cancer.

Meanwhile, a variety of advanced imaging modalities have emerged to probe soft tissues in three dimensions, offering deeper insights into the tumor microstructures [16]. Fluorescence-based techniques—of which the most promising is light-sheet microscopy operate at optical wavelengths and provide cellular resolution with cell-specific contrast [17]. However, these methods typically require extensive non-reversible chemical sample preparation, i. e. tissue clearing and fluorescent staining [18]. By contrast, micro-CT can accommodate larger samples, is fast, and does not necessitate additional chemical treatments, enabling non-destructive visualisation of the tumor micro-architecture at high fidelity, though it lacks cell-specific contrast and resolution [19].

Micro-CT has the potential to move beyond being a sole research tool by providing clinical benefits. It has shown the ability to support histology of FTCs by providing a much-needed 3D insight about the presence of vascular or capsular invasions [7] [20]. Micro-CT has also been utilized in propagating 2D spatial transcriptomics to 3D volume [21, 22], and prognostics of prostate cancer, where it outperforms traditional histology by using 3D information [23]. Micro-CT integrates seamlessly into clinical workflows and does not require any sample preprocessing other than formalin-fixed paraffin-embedded (FFPE) block preparation.

Since biological processes influence tissue texture, inverse-modelling of image texture can be used to infer molecular profiles, stratify risk, and produce other diagnostic read-outs [24]. Classical radiomics—built on hundreds of hand-crafted intensity, shape, and texture descriptors—has already connected such imaging signatures to actionable genetic alterations while remaining transparent and interpretable [25]. Deep-learning pipelines learn richer, task-specific representations, but they typically require large, well-annotated datasets and their latent features can be opaque to pathologists [26]. Foundation vision models trained on millions of heterogeneous images offer a more adaptable backbone that generalises across tumour types, staining protocols, and imaging modalities, holding particular promise for computational pathology [27].

Combining computer vision models with advanced imaging techniques, such as micro-CT paves the way for identifying surrogate imaging biomarkers, improving 3D tissue characterization, and driving progress in precision diagnostics for cancer research [28]. However, the specificity and sensitivity of such models must be rigorously validated before any clinical adoption—unless their role is limited to screening, where different performance thresholds may be acceptable.

Clinical benefits of radiomics have been demonstrated across a wide range of tasks and imaging modalities. It has been applied to predict lymph node metastasis in PTC using multiple imaging modalities, including ultrasound [29], CT [30], and MRI [31]. Beyond PTC, MRI radiomics has proven useful for detecting TERT promoter mutations in glioblastoma [32], as well as IDH mutation status in glioma [33]. PET/CT radiomics has facilitated the identification of EGFR mutation subtypes in lung adenocarcinoma [34]. Additional research has demonstrated that cardiac cine MRI can help stratify hypertrophic cardiomyopathy patients by their risk of heart failure [35], and that CT-based radiomics can predict KRAS mutations in colorectal cancer [36].

Micro-CT radiomics contributes to the evolution of quantitative pathology, where diagnostic decisions are increasingly guided by reproducible, data-driven metrics. Through high-resolution, three-dimensional imaging, this technique reduces sampling errors and provides structural detail beyond the abilities of standard two-dimensional slides. We applied micro-CT radiomics to classify thyroid tumors across several clinically relevant tasks on a large cohort of thyroid tumors. First, we reliably distinguished tumor tissue from non-neoplastic thyroid parenchyma. Subsequently, radiomic features enabled the classification of PTC and follicular thyroid neoplasms (FTN), encompassing both follicular thyroid carcinomas and adenomas. Finally, the features were used to predict BRAF V600E mutation status.

Through the SHapley Additive exPlanations (SHAP) of the classification model [37], we uncovered the key textural patterns driving each classification. We also observed preliminary associations between specific radiomic features and TERT promoter mutations. Finally, no radiomic features reliably predicted RAS Q61R mutation status or risk of relapse in thyroid tumors.

The method presented here shows strong potential to serve as an efficient, cost-effective, and multi-channel screening approach for large tumor volumes. By capturing spatial heterogeneity across the entire lesion, it may contribute to a deeper understanding of thyroid tumor biology and support more personalized patient management.

## 2 Results and discussion

### 2.1 Tissue type

We first evaluated the model’s ability to distinguish non-neoplastic (n=334) from neoplastic (n=403) thyroid tissue. As shown in the Uniform Manifold Approximation and Projection (UMAP) projection Fig. 1(a) [38], radiomic features form two separated clusters, indicating that the model captures biologically meaningful, class-discriminative patterns. Receiver operating characteristic (ROC) analysis across the validation folds in a 5-fold cross-validation yielded an area under the curve (AUC) of 0.87 ± 0.03 Fig. 1(b), confirming robust generalisation. The predicted probability distributions Fig. 1(c) exhibit distinct bimodality, with minor overlap between classes due to biological gray zones or a mixed presence of neoplastic and non-neoplastic tissue in some tissues. Additional performance metrics and dataset details are provided in table 5. Together, these results demonstrate that our radiomics model reliably identifies the textural signatures that differentiate neoplastic from non-neoplastic tissue, laying a solid foundation for subsequent histopathological interpretation.

**Figure 1.**
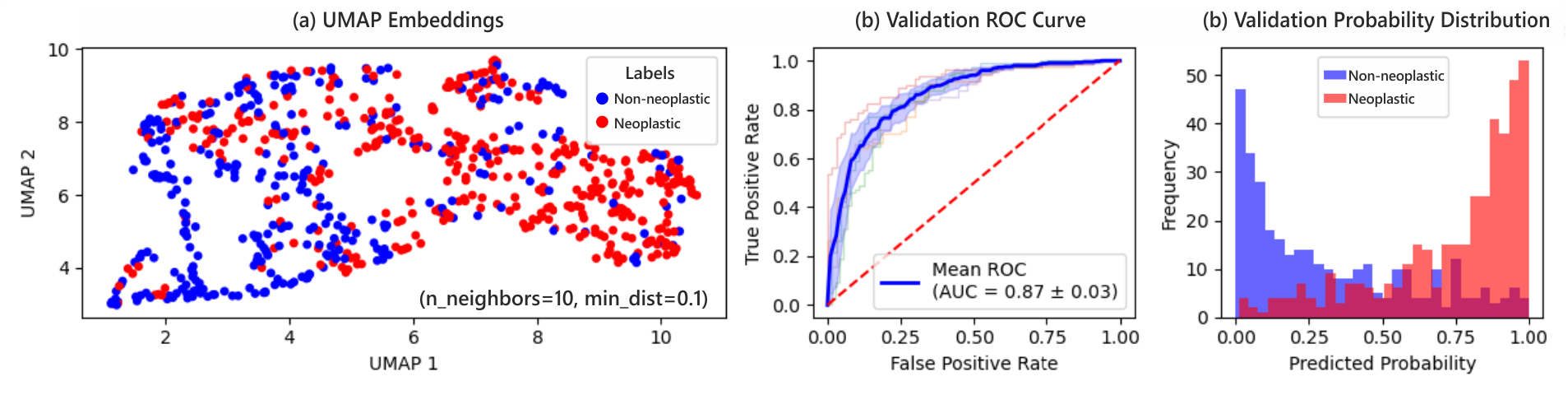
Tissue type classification, (a) the UMAP of the radiomics embeddings. (b) the ROC curve, and (c) the aggregated probability distribution across validation folds.

### 2.2 Tumor type

Radiomics embeddings showed strong performance for FTN (n=109) and PTC (n=96), which together account for up to 95% of thyroid neoplasms. Yet, they lacked discriminative power for poorly differentiated and oncocytic carcinomas. The dataset details of the classification metrics are given in table 5.

The UMAP visualization in Fig. 2(a) reveals partially separable feature representations between FTN and PTC. Nonetheless, the classifier achieved a validation AUC of 0.84 ± 0.05 in Fig. 2(b), indicating robust generalizability across validation folds. Furthermore, the model’s probability distribution in Figure Fig. 2(d) shows a distinct bimodal pattern, supporting the model’s ability to differentiate the two tumor groups despite some overlap.

**Figure 2.**
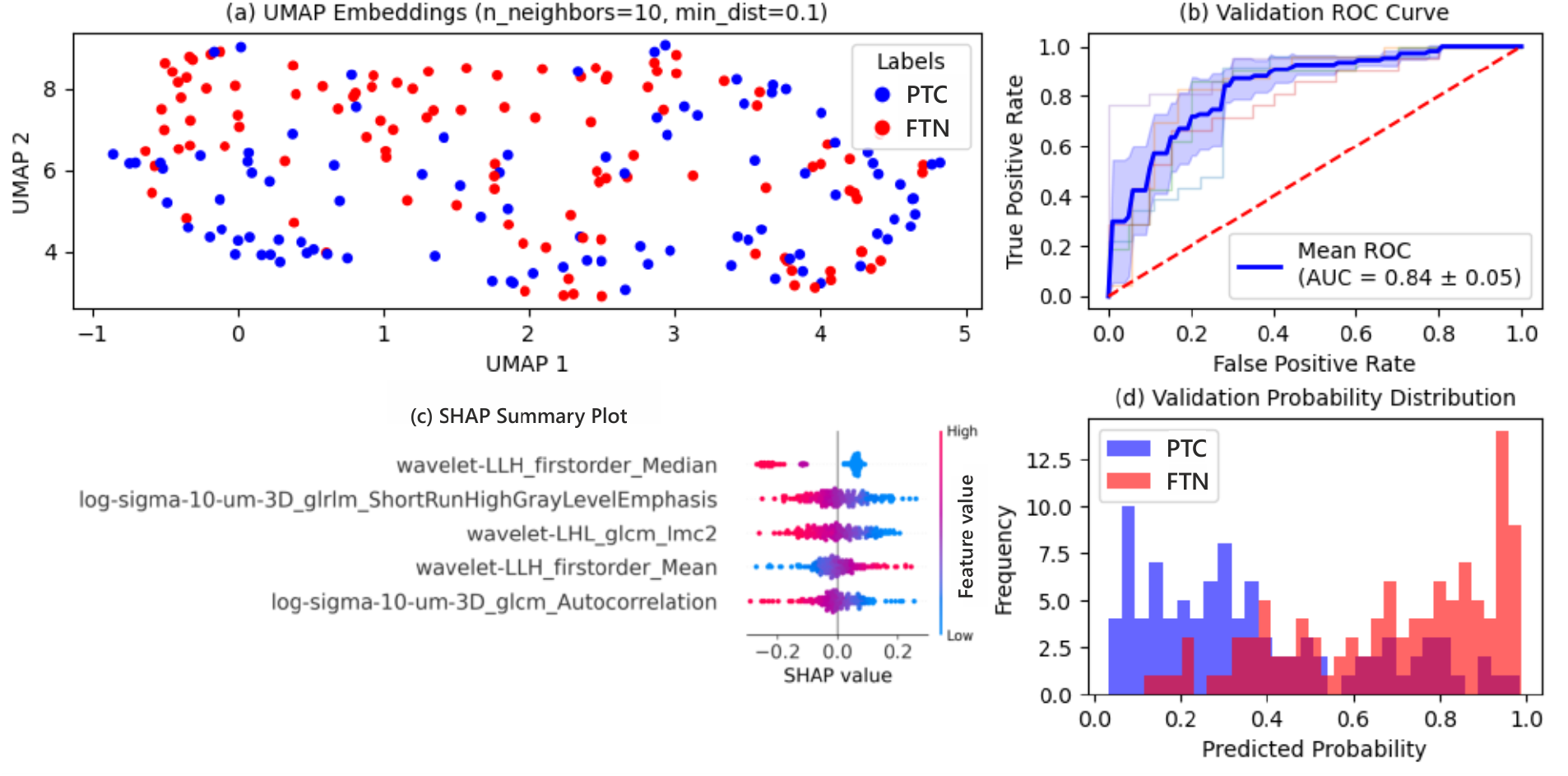
Model performance PTC and FTN classification. (a) UMAP visualization of radiomics embeddings shows partial class separation between PTC (blue) and FTN (red). (b) ROC curve on validation data with a mean AUC of 0.84 ± 0.05. (c) SHAP summary plot highlights top 5 most impactful radiomic features. (d) Histogram of predicted probabilities shows a bimodal, distribution on the validation set.

SHAP analysis in Fig. 2(c) shows that first-order statistics, namely the median and mean from the wavelet LLH band (see Fig. 9 for an example), contribute significantly to the classification. Higher median and lower mean values—a gray level distribution with numerous moderate-intensity transitions—favor PTC, whereas the inverse pattern, lower median and higher mean–Fewer transitions but sharper in magnitude–indicates FTN. Laplacian of Gaussian (LoG)-GLRLM-short run high gray level emphasis also correlates positively with PTC, indicating the presence of many short runs of grey values corresponding to 10 µm structures in the original image Fig. 9. Additionally, from the same wavelet band a higher GLCM-Imc2, and from LoG, GLCM-autocorrelation correlate positively with PTC. This implies that the transitions are regular and repetitive; The indicated patterns are in-line with the PTC (**) and FTC (*) punches shown in Fig. 7(b), and the expected follicular growth pattern in FTCs.

The proposed method enables accurate classification of PTC and FTN and may serve as an objective, quantitative tool for pathologists to quantify growth patterns and tumor textural activity. By analyzing the entire tumor volume, it offers an efficient approach for comprehensive subtype classification. The model’s tendency to favor FTN–characterized by fewer but more pronounced intensity transitions–aligns with the follicular growth pattern, where iodine-rich follicles produce bright, sharply defined structures in X-ray images.

### 2.3 Follicular variant of papillary thyroid carcinomas

Clinically, FVPTC is recognised as an intermediate entity between PTC and FTC with nuclear features of PTC and invasive growth pattern of FTC [39]. Our findings provide radiomics-based evidence supporting this intermediate status. The radiomics classification between FVPTC and PTC achieved a modest 5-fold cross-validation AUC of 0.67 ± 0.12, indicating their textural affinity. Classification between FVPTC and FTN yielded a slightly higher AUC of 0.71 ± 0.07. Additionally, classifying FVPTC against a combined group of PTC and FTN resulted in a validation AUC of 0.67 ± 0.04 (see table 1).

**Table 1:** Radiomics-based classification performance (5-fold cross validation AUC) of FVPTC against PTC and FTN.

| Comparison | AUC (mean ± SD) |
| --- | --- |
| FVPTC (n=53) vs. PTC (n=96) | $0.67 \pm 0.12$ |
| FVPTC (n=53) vs. FTN (n=109) | $0.71 \pm 0.07$ |
| FVPTC (n=53) vs. PTC + FTN (combined, n=202) | $0.67 \pm 0.04$ |

To further investigate the imaging phenotype of FVPTC, we projected radiomic features into two dimensions using UMAP (Fig. 3). FVPTC samples appeared broadly distributed, yet showed a consistent spatial overlap with PTC rather than forming distinct clusters.

**Figure 3.**
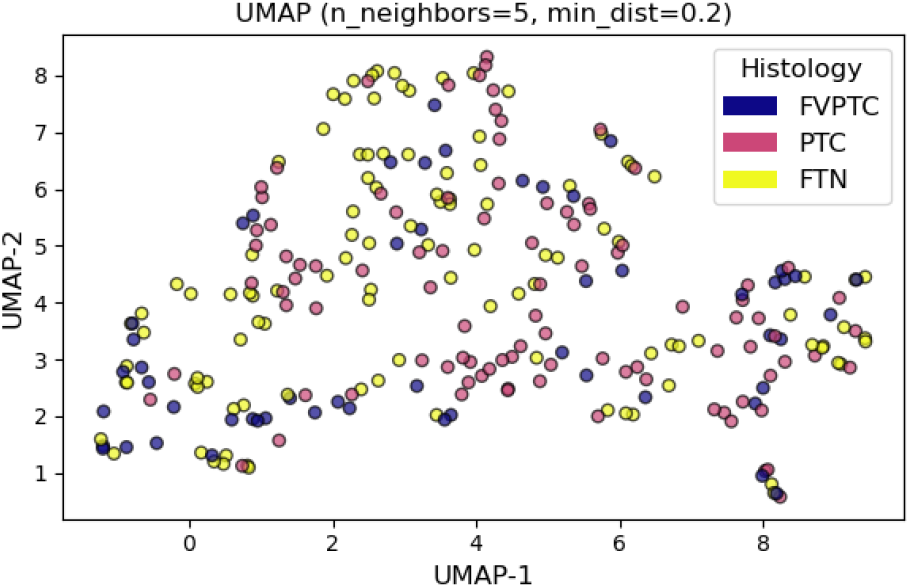
UMAP projection of radiomics features extracted from thyroid tumors. Each point represents a tumor sample, color-coded by histological subtype: FVPTC (blue), PTC (pink), and FTN (yellow). The projection reveals partial overlap between FVPTC and both PTC and FTN, with FVPTC clustering more closely with PTC.

These results confirm that FVPTC exhibits radiomics characteristics intermediate between PTC and FTN, albeit it might appear closer to PTC. This does not include nuclear features of PTC, since nuclear information is not resolved by micro-CT. This closer affinity reflect textural activity as detailed in section 2.2. The modest classification scores reflect the intrinsic diagnostic ambiguity of FVPTC, which is also known to be associated with high intra- and inter-observer variability [5]. In this context, a model showing low-confidence classification between PTC and FTN may serve as an indicator of FVPTC, providing a more objective assessment in equivocal cases. This model can contribute to a more reproducible and data-driven evaluation of this diagnostically challenging subtype.

### 2.4 BRAF V600E status

The BRAF gene encodes a kinase involved in the MAPK signaling pathway, which regulates cell proliferation and survival. Mutations in BRAF, particularly V600E, result in constitutive pathway activation and are frequently implicated in tumorigenesis. BRAF mutation is highly prevalent (89%) in aggressive classical PTC [40], making it a key prognostic marker. In contrast, BRAF mutations are absent in FTN [10], therefore, FTNs were excluded from the dataset of this classification task. The dataset and details of the classification metrics are given in table 5. While BRAF mutations are generally homogeneous [41], intratumoral heterogeneity—particularly of BRAF status in multifocal [13], and polyclonal PTCS has been reported [15], highlighting the potential value of 3D imaging modalities like micro-CT for mutation status classification.

The classification performance of the proposed model for predicting BRAF V600E mutation status is summarized in Fig. 4. The UMAP projection Fig. 4(a) demonstrates a degree of class separability between BRAF-positive (n=79) and BRAF-negative (n=166) samples in the radiomics feature space, indicating that the extracted radiomic features capture relevant discriminatory information. ROC analysis Fig. 4(b) yields a mean AUC of 0.77 ± 0.06 in 5-fold cross validation, suggesting that the model achieves moderate discrimination between two classes. The relatively small standard deviation reflects consistent performance across validation folds. The predicted probability distributions Fig. 4(d) show a reasonable separation between classes. Nonetheless, overlap exists, which could reflect borderline cases or noise in the data. The model’s potential to screen the full tumor volume for BRAF V600E and map intratumoral heterogeneity is evident. However, improvements are still needed to increase specificity, such that the method may ultimately support reliable stand-alone mutation analysis.

**Figure 4.**
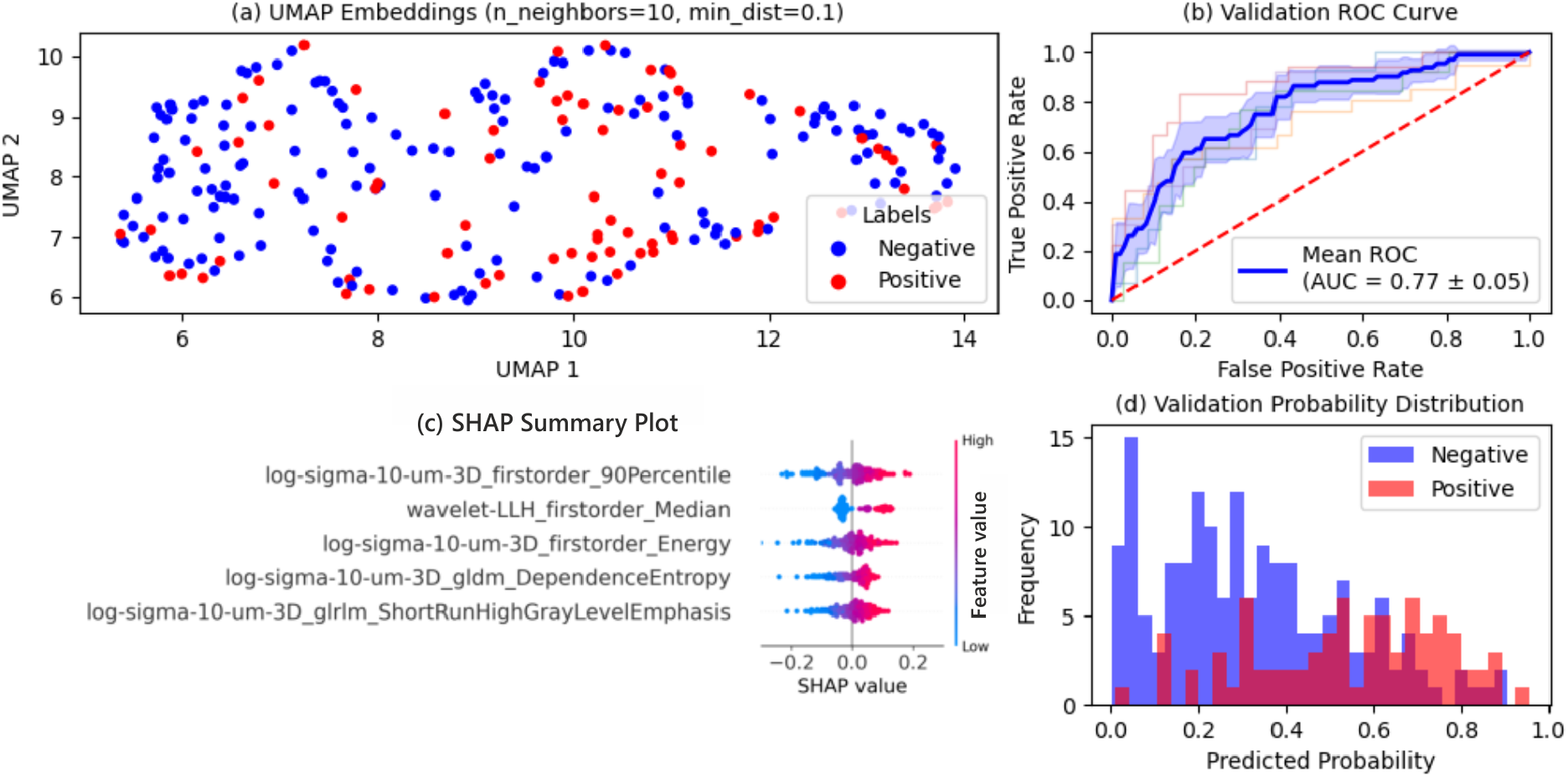
(a) UMAP visualization of feature embeddings shows separation between wild-type (blue) and mutated (red) classes. (b) ROC curve for the validation set with an average AUC of 0.77 ± 0.05, indicating good discriminatory ability. (c) SHAP summary plot highlighting the top contributing radiomic features; color indicates feature value (blue: low, red: high) and x-axis shows SHAP value impact. (d) Histogram of predicted probabilities in the validation set, showing class separation in model outputs.

Fig. 4(c) shows the SHAP summary for the classification of BRAF mutation status. The most informative radiomics features consistently highlight fine-scale intensity and texture variations, particularly at the 10 µm level. Features derived from the LoG filter class—such as firstorder 90percentile, Energy, GLRLM short run high gray level emphasis, as well as wavelet firstorder median suggest that BRAF-mutated tumors exhibit more intense and frequent textural activity at this microscopic scale. The GLDM dependence entropy feature further supports this by capturing the elevated spatial heterogeneity and irregularity of mutated tumors. An example of a BRAF-positive tumor is shown in Fig. 5(a) with higher textural activity and heterogeneity consistent with the SHAP analysis in section 2.4. The classifier predicts 84% and 36% BRAF mutation probability for Fig. 5(a), and (b), respectively. Below each punch, the immunohistochemistry image for BRAF V600E is shown.

**Figure 5.**
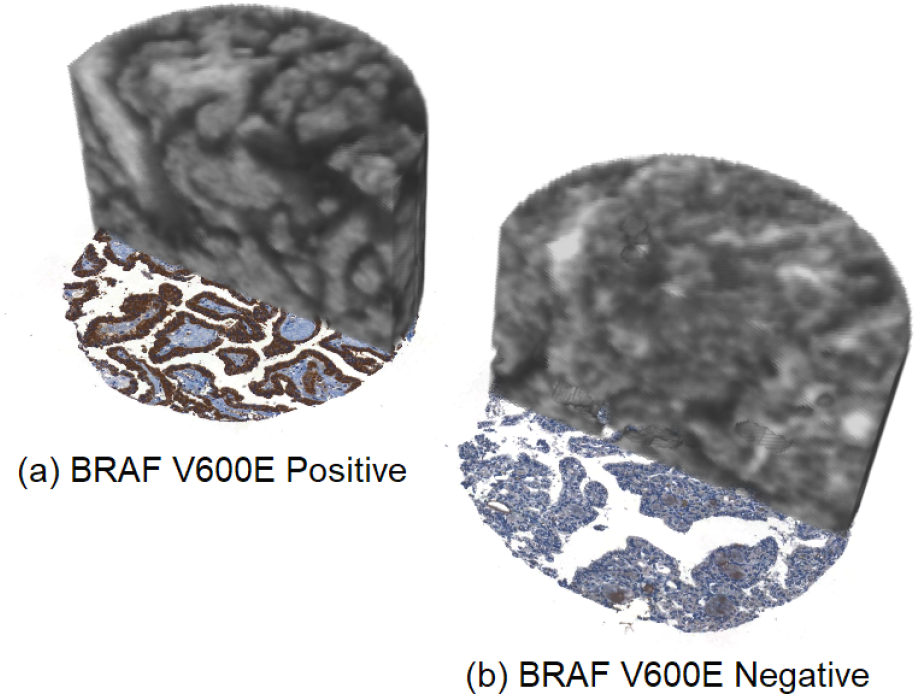
Two punches, (a) BRAF positive, and (b) BRAF negative, their corresponding BRAF immunohistochemistry image is shown at the bottom of punch. Each punch is 600µm in diameter.

### 2.5 TERT status

Since their discovery, the TERT promoter mutations have emerged as key prognostic genetic alterations in thyroid cancer. These promotor mutations result in de novo ETS transcription factor binding motifs, leading to increased TERT expression and cellular immortalization. The prevalence of TERT mutation in benign tumors is zero, while it increases to 11.3% in PTC, 17.% in FTC, 43.2% in PDTC, and 40.1% in ATC. Their enrichment in aggressive subtypes underscores their association with adverse clinical outcomes, including recurrence and mortality [42]. These findings position TERT promoter mutations as promising biomarkers for risk stratification and potentially therapeutic targeting in thyroid cancer. Motivated by its significance, we used the radiomics embedding for TERT classification on our limited available dataset with 8 mutated and 108 wild-type samples (see table 5).

Fig. 6(a) presents the validation ROC curve with a mean AUC of 0.76 ± 0.15, suggesting moderate discriminative performance but with notable variability across validation folds. Fig. 6(c) displays the distribution of predicted probabilities. Although mutated samples are consistently scored on the higher end of the predicted probabilities, the model is not confident as all predictions are below 0.5 due to the heavily biased and small dataset.

**Figure 6.**
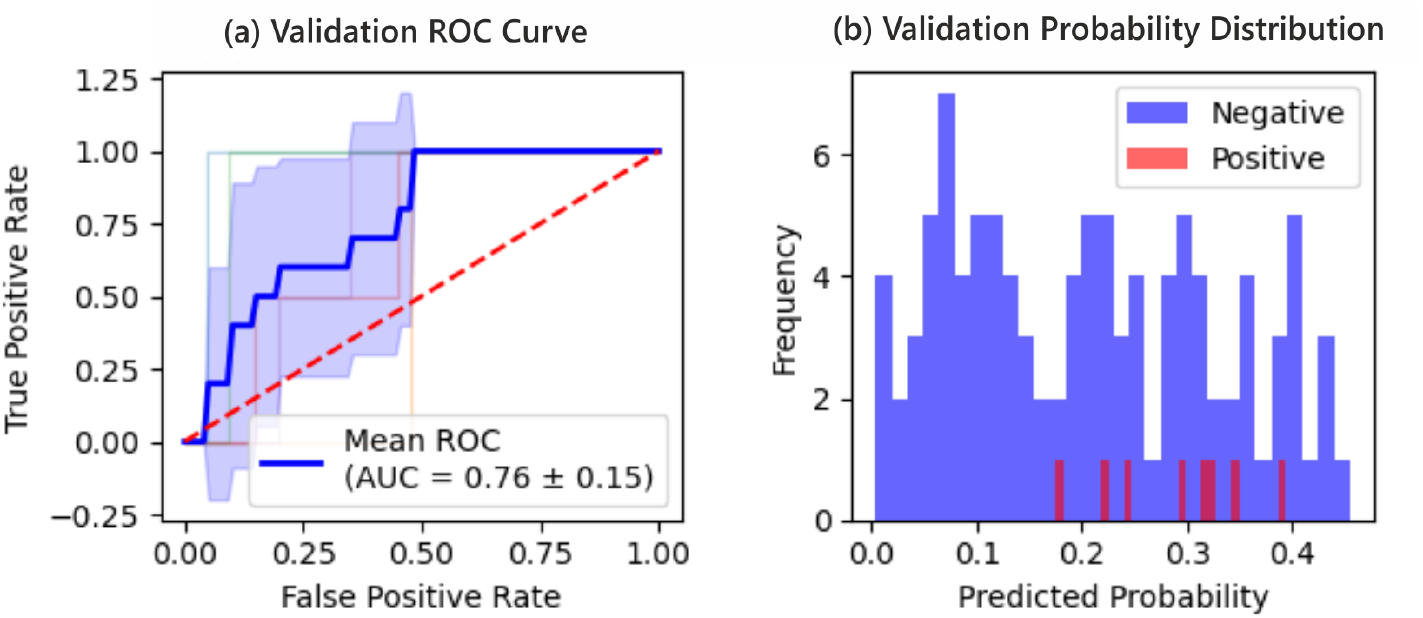
Evaluation of TERT mutation status classification. (a) Validation ROC curve showing a mean AUC of 0.72 ± 0.16. (b) Distribution of predicted probabilities.

**Figure 7.**
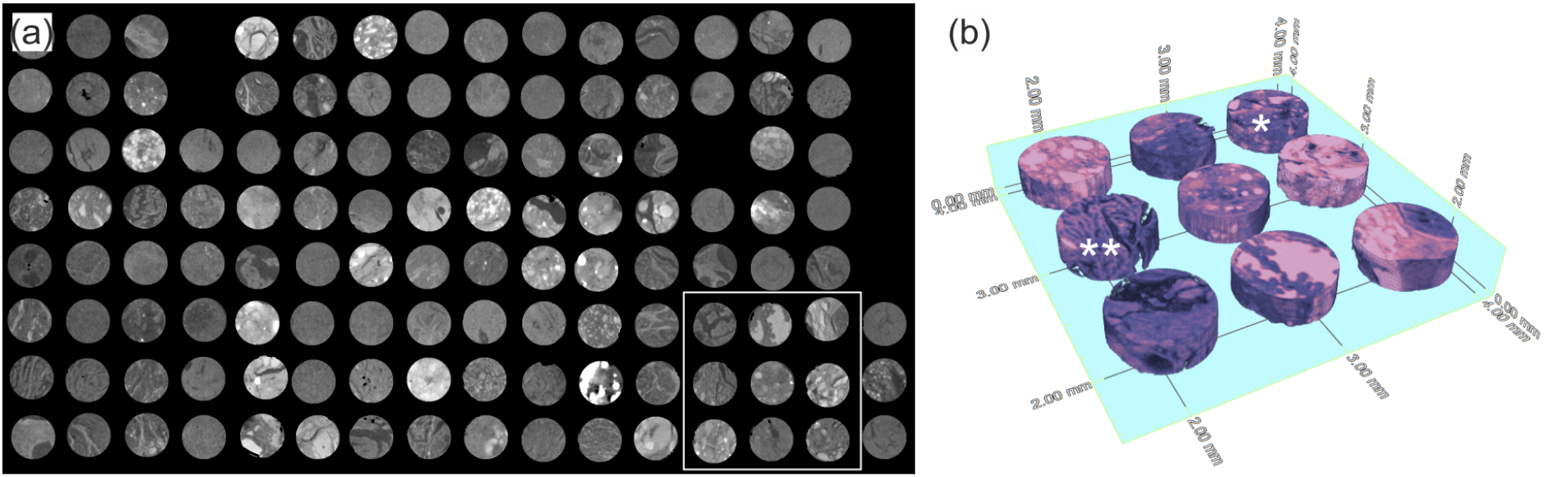
(a) A slice of an ngTMA batch with 120 punches, (b) A false color 3D rendering of a 3by3 grid of ngTMA area indicated by the white rectangle in (a). The punches annotated by * and ** are FTC and PTC, respectively.

To better understand the underlying signal, we applied the non-parametric Mann–Whitney U test to each radiomic feature, comparing TERT-mutated and wild-type tumors. A subset of features showed statistically significant differences between groups (see table 2), suggesting that certain textural traits may be enriched in TERT-mutated tumors. These findings, while preliminary, support the hypothesis that generalizable radiomic markers of TERT mutation may exist. Further research and validation of this cost-effective, fast, and high-throughput 3D screening of molecular alterations are required.

**Table 2:** Mann–Whitney U test results comparing radiomic features between TERT-mutated and wild-type tumors.

| Feature | U-statistic | p-value |
| --- | --- | --- |
| Squareroot-GLSZM-high gray level zone emphasis | 217 | 0.0246 |
| Wavelet-HLH-GLSZM-zone percentage | 570 | 0.0721 |
| Original-firstorder-skewness | 262 | 0.0885 |
| Original-firstorder-kurtosis | 264 | 0.0930 |
| Wavelet-HLL-GLDM-dependence variance | 265 | 0.0953 |

### 2.6 RAS Q61R status

Here, using a dataset of 308 wild-type and 46 RAS Q61R mutated tumors, we found that there exists no generalizable radiomics signature from micro-CT images at this resolution scale (10 µm) that encodes the appearance of RAS Q61R mutated tumors (table 5). This is not surprising, as other RAS mutations—not detected by the mutation-specific immunohistochemistry—can lead to similar morphologies and molecular alterations [43].

### 2.7 Relapse Risk

The evaluation of the relapse prediction model using 128 disease-free survival and 95 relapsed cases yielded performance comparable to random classification (table 5), indicating no meaningful association between textural features and relapse risk. These results suggest that, at the current imaging resolution, textural markers for relapse do not exist. Relapse prediction is known to be a multifactorial problem, with a strong influence of tumor stage and age of the patient, besides the intrinsic tumor characteristics. The absence of meaningful texture-based predictors for relapse prediction underscores the complexity of these clinical outcomes, which likely depend on biological factors beyond microstructural properties alone.

## 3 Conclusion

The integration of advanced machine vision models with micro-CT—offers new opportunities to capture complex morphological and textural patterns. In this study, we demonstrated that radiomic analysis of 3D micro-CT images enables reliable classification of FTNs and PTCs, and BRAF V600E mutation. Our preliminary findings suggest that there may exist radiomics biomarkers for TERT promoter mutations status. These observations are based on a limited sample size and require further validation. Importantly, the 3D nature of the method allows capturing and mapping the intratumoral heterogeneity. The micro-CT radiomics presented here has the potential to evolve into an efficient, cost-effective, and multichannel screening tool for large tumor volumes—enabling systematic assessment of spatial heterogeneity and contributing to more personalized, biology-informed patient management based on quantitative pathology. Where feature values may guide and help pathologists beyond a binary classification.

## 4 Methods

### 4.1 Dataset

The next-generation tissue microarrays (ngTMA) represent a key enabler for scaling this technology [44]. By imaging multiple tissue cores under uniform conditions, ngTMAs reduce scan time, increase throughput, and ensure consistency in radiomics feature extraction, addressing a major bottleneck in the adoption of 3D imaging modalities for data-driven diagnostics tasks.

Fig. 7(b) illustrates a 3D rendering of a micro-CT scan depicting a 3×3 grid subvolume from a ngTMA, artificially colored to mimic hematoxylin and eosin staining. Each punch represents an individual specimen, with typical ngTMA blocks containing approximately 200 punches (see Fig. 7(a)). Our study analyzed four ngTMA blocks, resulting in a dataset comprising 334 non-neoplastic and 403 neoplastic tissue samples. Most patients provided paired punches—one from non-neoplastic peripheral tissue and one from neoplastic tissue. The overall cohort included 418 patients, though effective sample sizes varied according to specific classification tasks and the tumors data availability, as detailed in table 5.

### 4.2 X-ray micro-CT

X-ray micro-CT was done using a commercial device – EasyTom XL Ultra (Rx Solutions, Chavanod, France). The scanner features a Hamamatsu reflection target microfocus X-ray source L10801. The tube was operated at 90kVp, with 110µA, resulting in the smallest possible tube spot size at 5µm. The detector used in this study is a 1880 × 1494 pixel array of flat panel with a physical pixel size of 127µm and 16-bit dynamic range.

The scintillator is a 700µm thick columnar CsI. The exposure time was set to 0.5s to utilize the full dynamic range of the detector, and for each projection, 30 frames were averaged to increase the signal-to-noise-ratio. Total number of projections over 360 degrees per scan is 1440, resulting in 6 hours of theoretical scan time.

The ngTMA blocks were placed vertically on the scanner to maximize the contrast-to-noise-ratio and the scan geometry was designed to maximize the photon economy at the desired voxel size rather than edge enhancement [19]. This compromise was made due to hardware limits and time constraints. The SDD was 667.35mm and the SOD was 26.27mm, resulting in a voxel size of 5.00µm and an effective area coverage of 9.4mm by 7.5mm.

Given the 600µm diameter of each punch and the spacing between the punches, during each scan approximately an array of 7 by 5 punches could be scanned. Therefore, each ngTMA block is scanned in multiple individual scans. Then these scans are stitched together to form the complete array. Consequently, depending on the number of punches in an array, and the spacing between the punches, the net scan time of the ngTMAs can vary.

Then, each individual scan is reconstructed using the commercial reconstruction software, Xact (RX Solutions, Chavanod, France). Where a set of trivial image processing step are applied, first a ring filter correction is applied, followed by a Paganin filter with a 0.25f_Ny_ cut of frequency at − 6dB [45]. Then a standard filter back projection algorithm [46], is applied to reconstruct the volume. Which is then mapped to a fixed unified gray value contrast window for all the individual scans.

### 4.3 Image processing

#### 4.3.1 Segmentation

The segmentation process begins by computing a mean intensity projection along the z-axis, transforming the 3D ngTMA volumes into a single 2D image. To reduce noise sensitivity inherent to the subsequent steps, a Gaussian smoothing filter with a kernel size of 5 pixels is applied. Next, Hue’s circle transform from OpenCV is utilized on the smoothed image, using a predefined circle radius of 65 pixels. Detected circles are aligned against a reference grid generated based on the image dimensions and known punches layout. Each detected circle is assigned a gray value linked to specific grid coordinates.

Following circle detection, the circles are duplicated along the depth, creating cylindrical volumes encapsulating each punch. Regions corresponding to air inclusions (identified by zero gray value pixels from the embedding artifacts) are excluded from these cylinders. These exclusion zones are then expanded through morphological dilation using a spherical kernel of 2 pixels, effectively removing embedding artifacts and ensuring their artifacts do not propagate by the downstream image processing steps into the feature extraction process.

#### 4.3.2 Filter classes

In the Pyradiomics library, if the filter class is activated, it is calculated on the original image and then rescaled [47]. However, in the case of ngTMAs, it does not work properly. In this scenario, the air embedding artifacts give rise to false signals that tend to dominate in some filter classes (e. g. gradient filter), because of the abrupt change in the gray value at the air inclusions. These dominating signals corrupt the subsequent normalization process, and considering the image discretization that comes naturally in radiomics, all the gray values collapse in one or two bins. Consequently, we need to apply the filters and normalize them manually and feed them as original images to the library. We applied the following filter classes using “pyradiomics.imageoperations”. **(i)** Laplacian of Gaussian (*σ* = 10*µm*), **(ii)** wavelet (start_level=0, level=1, wavelet=“haar”), except HHH, **(iii)** logarithm, **(iv)** square Root, **(v)** gradient, **(vi)** exponential, **(vii)** square.

The directional channels in the wavelet decomposition are not meaningful due to the isotropic nature of both tissues and voxel sizes. As a result, automatically only one direction is retained for each feature across the three directional channels after redundancy reduction (see section 4.5.3). In the SHAP analysis described in section 4.6.2, any directional information from wavelet channels should be disregarded. For instance, wavelet-LLH, HLL, and LHL are considered equivalent.

#### 4.3.3 Normalization

Based on the visual observation of the filtered images, a histogram window is selected such that it does not saturate the upper histogram and does not cut off gray values at the lower end of the histogram. For each filter class, this window is unified and does not change from one scan to another. Normalized images are then discretized to 8 gray levels to avoid sparse matrices.

### 4.4 Feature extraction

Feature extraction is done using PyRadiomics library [47] with 8 gray levels. the activated feature classes are **(i)** First order, **(ii)** gray level co-occurrence matrix (GLCM), **(iii)** gray level run length matrix (GLRLM), **(iv)** gray level dependencies matrix (GLDM), **(v)** gray level size zone matrix (GLSZM), **(vi)** neighbouring gray tone difference matrix (NGTDM). Shape features are not extracted since the shape of all the cores is a cylinder. This feature class setting produces 93 features per filter class, resulting in 1302 features.

### 4.5 Feature Processing

Many of 1302 features are not robust to variations in the imaging protocol, and many of them are rudimentary. Last but not least, the variations in the parrafin properties and its thickness result in batch bias. Therefore, a set of feature processing steps are necessary before using the embeddings for classification tasks.

#### 4.5.1 Non-reproducible features

While radiomic features provide valuable diagnostic signals, their interpretability and generalizability across cohorts and scanner platforms remain active areas of investigation [48]. We took a rigorous approach to ensure the reproducibility of the features. We evaluated the intraclass correlation coefficient (ICC) between two distinct micro-CT scans of a single ngTMA block containing 193 punches. The first scan utilized previously described standard parameters (as detailed in Section section 4.2), while the second scan involved positioning the ngTMA block horizontally, significantly altering imaging artifacts and quality metrics. Apart from the mounting mode of the sample, scanning parameters were selected such that they are widely different.

The first scan is from the dataset acquired through the protocol described in section 4.2, the second scan featured modified geometry: source-to-detector distance (SDD) of 856.14 mm and source-to-object distance (SOD) of 33.72 mm, resulting in a voxel size of 5.00 µm. The exposure time per projection was 0.67 s, with 40-frame averaging across 2912 projections over a full 360-degree rotation. Additionally, we virtually expanded detector coverage by horizontally offsetting and stitching two projections per angle, effectively doubling detector width and achieving an expanded field of view. This method resulted in a theoretical total scan duration of approximately 43 hours and 8 minutes for a full ngTMA block.

Images from both scans underwent identical reconstruction and embedding procedures, consistent with the methodologies outlined in Sections section 4.3 and section 4.4, to ensure validity for ICC analysis. Next, intraclass correlation was calculated between the two sets of features, using the “intraclass_cor” function from the Penguin library in Python. Features with an ICC smaller than 0.75 and a p-value smaller than 0.05 were discarded as non-reproducible.

#### 4.5.2 Batch correction

Batch effects in ngTMA radiomics datasets can arise from variations in chemical properties of paraffin wax, subtle differences in scanning geometry, or hardware instabilities during image acquisition. These batch-related variations introduce non-biological discrepancies among extracted radiomic features.

Prior to correction, clustering of the six ngTMA batches is clearly observed in the UMAP visualization of the radiomic feature space, as demonstrated in Fig. 8(a). First a sign preserving log-transform is performed, then following batch correction using ComBat [49], from the pyCombat package [50], the batch-related clustering is effectively mitigated, yielding biologically-driven feature distribution as illustrated in Fig. 8(b). ComBat was chosen due to it is previously demonstrated effectiveness in radiomics applications [51], providing robust correction of batch effects.

**Figure 8.**
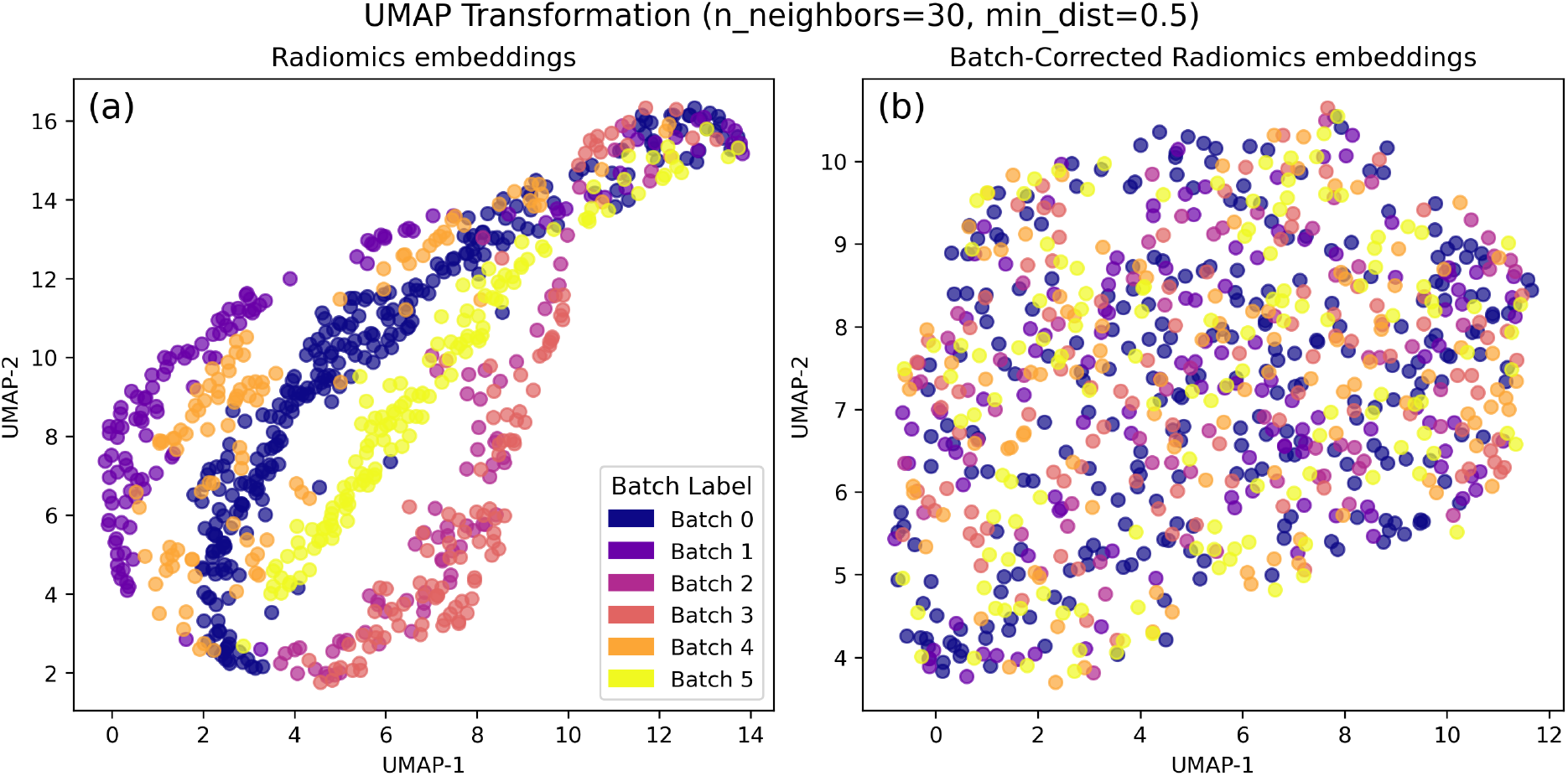
UMAP of dataset (a) before and (b) after batch correction.

**Figure 9.**
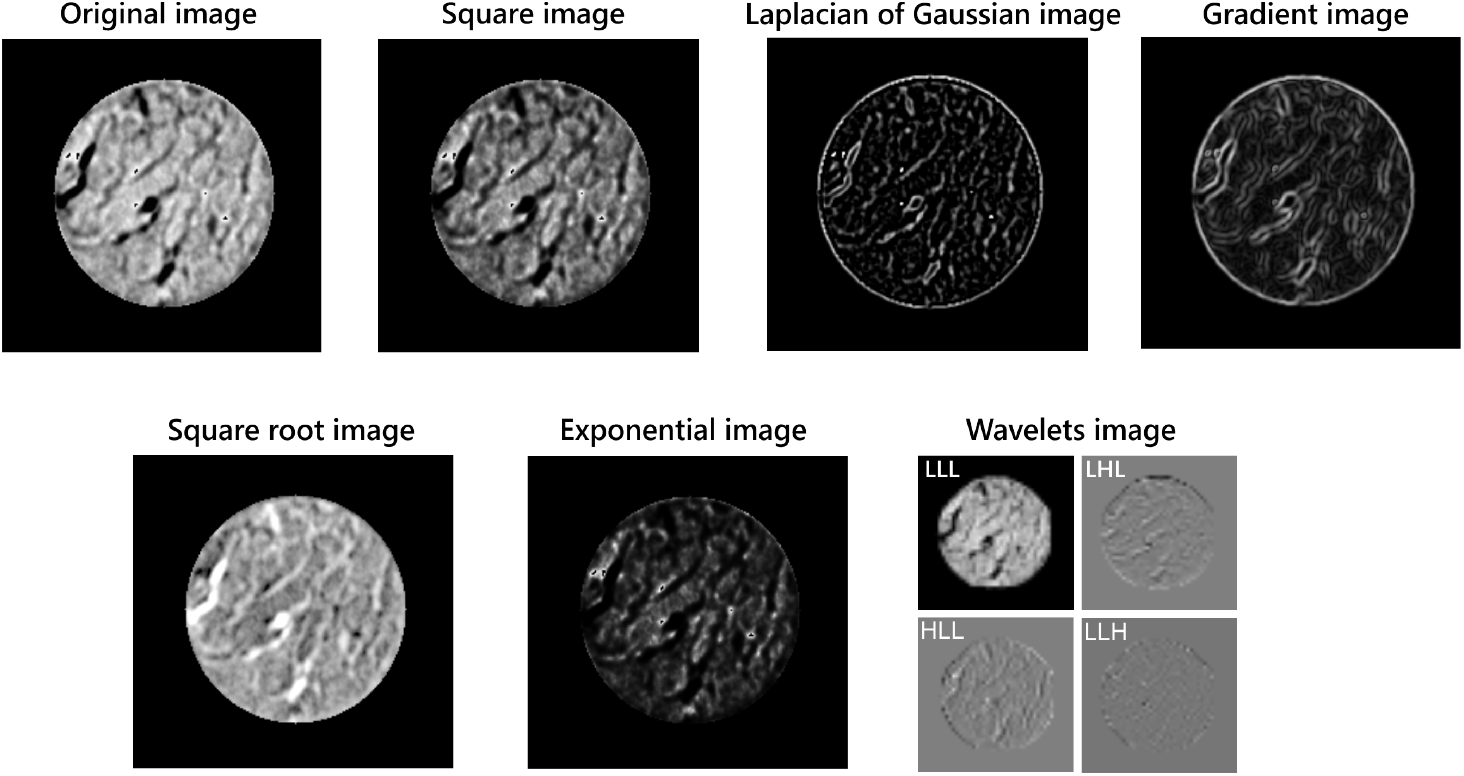
The images demonstrate various filter classes used in radiomic feature extraction. Each image highlights a different aspect of the tissue.

#### 4.5.3 Redundancy reduction

Following batch correction, feature redundancy is addressed by calculating Pearson correlation coefficients between all possible feature pairs using the SciPy library in Python. Feature pairs exhibiting a correlation coefficient greater than 0.95 and a p-value less than 0.05 are identified as highly redundant. From each redundant pair, one feature is randomly selected and discarded iteratively until all redundancy is resolved. We ended up with 151 features that are robust and informative. The resulting feature distribution by filter and feature class is given in table 3.

**Table 3:**
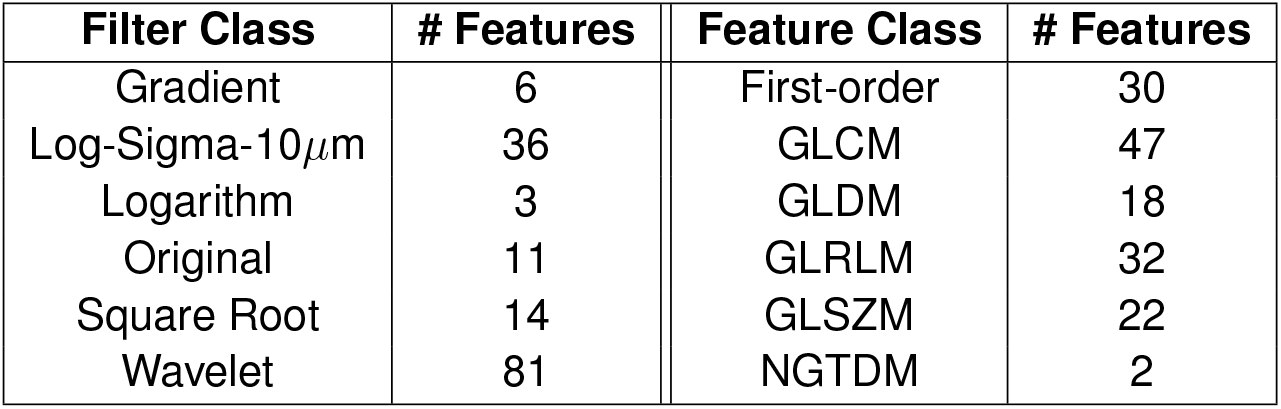
Distribution of features by filter and feature class.

### 4.6 Classifier

#### 4.6.1 Architucture and training

We trained a feed-forward neural network (Multilayer Perceptron, MLP) composed of two fully connected layers. We chose MLP because of its flexibility and its ability to capture non-linear relationships between features. The input to the model is a 151-dimensional radiomics embedding. The first layer projected the embedding into a hidden representation of dimension H (a hyperparameter), which was followed by a ReLU activation and a dropout layer to mitigate overfitting. The second layer was a fully connected output layer mapping the hidden features to the target classes. We trained the model using an adam optimiser by minimizing a standard cross-entropy loss with weight decay to regularize the model parameters. We used a learning rate scheduler for stable convergence. This architecture captures non-linear relationships among the radiomics features while remaining compact and straightforward to optimize.

The following hyperparameters were kept constant across all classification tasks: learning rate (0.001), weight decay (0.001), and learning rate scheduler gamma (0.95). Varying hyperparameters for each classifier are given in table 4.

**Table 4:** Classifier hyperparameters that vary across classification tasks. Shared parameters are listed in the section 4.6.1.

| Hyperparameter | Tissue type | Tumor type | BRAF V600E | TERT | RAS Q61R | Relapse |
| --- | --- | --- | --- | --- | --- | --- |
| Hidden Dim | 256 | 128 | 256 | 64 | 128 | 128 |
| Num Epochs | 10 | 5 | 5 | 2 | 4 | 5 |
| Class weights | [1, 1] | [1, 1] | [1, 2] | [1, 2] | [1, 2] | [1, 1] |
| Batch Size | 16 | 8 | 16 | 4 | 16 | 16 |
| Dropout rate | 0.6 | 0.2 | 0.3 | 0.7 | 0.3 | 0.3 |

**Table 5:** 5-fold cross validation training and validation metrics for each classification task.

| Task | Dataset | Training |  |  |  |  | Validation |  |  |  |  |
| --- | --- | --- | --- | --- | --- | --- | --- | --- | --- | --- | --- |
|  |  | Loss | AUROC | Precision | Recall | F1 | Loss | AUROC | Precision | Recall | F1 |
| Tissue type | Non-neoplastic: 334<br>Neoplastic: 403 | $0.34 \pm 0.01$ | $0.93 \pm 0.01$ | $0.85 \pm 0.01$ | $0.85 \pm 0.01$ | $0.85 \pm 0.01$ | $0.48 \pm 0.07$ | $0.87 \pm 0.03$ | $0.79 \pm 0.02$ | $0.79 \pm 0.02$ | $0.79 \pm 0.02$ |
| Tumor type | PTC: 96<br>FTN: 109 | $0.33 \pm 0.01$ | $0.95 \pm 0.01$ | $0.88 \pm 0.02$ | $0.88 \pm 0.02$ | $0.88 \pm 0.02$ | $0.52 \pm 0.05$ | $0.84 \pm 0.05$ | $0.75 \pm 0.07$ | $0.74 \pm 0.07$ | $0.74 \pm 0.07$ |
| BRAF V600E | Negative: 166<br>Positive: 79 | $0.41 \pm 0.02$ | $0.85 \pm 0.02$ | $0.85 \pm 0.02$ | $0.84 \pm 0.02$ | $0.85 \pm 0.02$ | $0.60 \pm 0.08$ | $0.77 \pm 0.05$ | $0.75 \pm 0.04$ | $0.73 \pm 0.05$ | $0.73 \pm 0.05$ |
| TERT | Negative: 108<br>Positive: 8 | $0.35 \pm 0.02$ | $0.83 \pm 0.04$ | $0.93 \pm 0.01$ | $0.53 \pm 0.04$ | $0.63 \pm 0.04$ | $0.38 \pm 0.03$ | $0.76 \pm 0.15$ | $0.92 \pm 0.04$ | $0.51 \pm 0.04$ | $0.62 \pm 0.04$ |
| RAS Q61R | Negative: 308<br>Positive: 46 | $0.43 \pm 0.01$ | $0.85 \pm 0.02$ | $0.89 \pm 0.01$ | $0.53 \pm 0.04$ | $0.59 \pm 0.04$ | $0.55 \pm 0.02$ | $0.54 \pm 0.09$ | $0.79 \pm 0.06$ | $0.45 \pm 0.09$ | $0.51 \pm 0.09$ |
| Relapse | Negative: 128<br>Positive: 95 | $0.57 \pm 0.01$ | $0.80 \pm 0.01$ | $0.72 \pm 0.02$ | $0.72 \pm 0.02$ | $0.70 \pm 0.02$ | $0.70 \pm 0.02$ | $0.59 \pm 0.04$ | $0.59 \pm 0.07$ | $0.59 \pm 0.06$ | $0.57 \pm 0.08$ |

#### 4.6.2 Classifier evaluation

All classifiers were evaluated using 5-fold cross-validation with the exception of TERT mutation status classifier, where we used a stratified 5-fold cross-validation due to the presence of only 8 positive samples. Performance metrics—including loss, accuracy-based scores, and area under the ROC AUC—were computed for both training and validation sets and are summarized in table 5. The accuracy besed metrics were calculated at the 0.5 probability threshold, except for the TERT mutation status classifier, where the threshold for positive class probability was lowered to 0.2. The probability distribution plots show the aggregated predicted class probabilities from all validation folds in the cross-validation analysis.

For 2D visualization of the dataset, we applied Uniform Manifold Approximation and Projection (UMAP) [38] to project the high-dimensional feature space into two dimensions. Probability distributions are reported as aggregated outputs across the cross-validation folds. To explain the model’s predictions, we used SHAP (SHapley Additive exPlanations) with the KernelExplainer method [37]. We defined a custom prediction function that applies the same scaling and model used during training. A set of 200 samples from the training data was used as the background for computing SHAP values. This allowed us to measure how much each feature contributed to the model’s output for each sample.

### 4.7 Immunohistochemistry

Immunohistochemical staining was performed on FFPE tissue sections. For BRAF V600E detection, the VE1 clone (Ventana Roche, 790-4855, RTU) was used. Antigen retrieval was carried out with Tris-EDTA buffer (pH 9.0) for 72 minutes at 99°C. Staining was performed using the Benchmark Ultra automated staining system (Ventana Medical Systems). For RAS Q61R detection, the SP174 clone (Abcam, ab227658, dilution 1:50) was applied. Antigen retrieval was performed with Tris-EDTA buffer (pH 9.0) for 40 minutes at 95°C. Staining was conducted on the Leica BOND III automated platform (Leica Biosystems). Notably, the antibody used detects Q61R mutations in HRAS, KRAS, and NRAS, while in PTC, NRAS and HRAS mutations are the most frequently observed.

### 4.8 TERT mutational Analysis

TERT promoter mutation analysis was performed as previously described [52]. Briefly, DNA was extracted from FFPE tissue using the DNeasy Blood and Tissue Kit (Qiagen, Hombrechtikon, Switzerland). A DNA fragment encompassing both mutational hotspot regions of the TERT promoter was amplified using the following primer pair: 5’-CACCCGTCCTGCCCCTTCACCTT-3’, and 5’-GGCTTCCCACGTGCGCAGCAGGA-3’. The PCR products were subsequently subjected to Sanger sequencing.

### 4.9 Data and Code Availability

The ngTMA images will be made available following completion of the peer review process. Radiomics embeddings (to be added post-review), corresponding labels, and all code used for image processing, feature extraction, and classification are publicly accessible at: GitHub Repository.

## 5 Ethical approval declarations

Tissue Biobank Bern (TBB) manages ngTMAs and patient information according to respective regulations and fulfills the Swiss Biobanking Platform requirements. The Institute of Tissue Medicine and Pathology (ITMP) of the University of Bern provided the ngTMAs and pseudonymized patient data from the TBB according to the protocol approved by the cantonal ethics commission (KEKBE 2018-01657).

## 6 Acknowledgements

This work was supported by the Strategic Focus Area Personalized Health and Related Technologies (PHRT) of the Eidgenössischen Technischen Hochschulen [Swiss Federal Institutes of Technology (ETH)] Domain, PHRT Pioneer Imaging Project “Toward Holistic Tissue Analyses: A Pioneer Imaging Project (PIP) for 3D Non-Invasive Histopathology of Thyroid Tumors for Precision Medicine” under Grant 2021-614.

## 8 Supplementary information

